# Overexpression of the MADS-box gene *CRM3* in the fern *Ceratopteris richardii* affects sporophyte development

**DOI:** 10.1101/2025.01.10.632496

**Authors:** Robin Schulz, Günter Theißen

**Affiliations:** Matthias Schleiden Institute / Genetics I, Friedrich Schiller University Jena, Philosophenweg 12, 07743 Jena, Germany

**Keywords:** CaMV 35S promoter, C-fern, MIKC-type gene

## Abstract

MADS-box genes encode MADS-domain transcription factors that play important roles in the development and evolution of land plants. Especially well-characterized is their role as organ identity genes during flower development of angiosperms. Even though the first MADS-box genes have been molecularly cloned from ferns almost 30 years ago, little is known about their function, mainly due to a lack of mutants. Also their quite broad expression domain, often comprising both the gametophytic and the sporophytic phase of the life cycle of ferns, provide few specific clues concerning gene function. However, as sister group of seed plants monilophytes (ferns and their allies such as horsetails) are of great interest in many attempts to understand both MADS-box gene and land plant evolution. Here we report, to the best of our knowledge, the first mutant phenotype based on transgenic overexpression of a fern MADS-box gene. Transgenic plants of the fern model system *Ceratopteris richardii* overexpressing the MIKC^C^-type MADS-box gene *CRM3* under control of the Cauliflower Mosaic Virus (CaMV) 35S promoter show a significant reduction of the length of sporophylls. This finding reveals that overexpression of *CRM3* affects sporophyll development in *C. richardii*, possibly by inhibiting cell proliferation or elongation.

## Introduction

MADS-box genes represent a eukaryote-specific family of genes encoding transcription factors, termed MADS-domain proteins (Gramzow et al., 2010). MADS-domain transcription factors play important roles in the development and evolution of plants and animals (Theißen et al., 1996, 2000, 2016). All MADS-domain proteins share a highly conserved, about 60 amino acids long DNA-binding domain, the MADS-domain (Käppel et al., 2023). MADS-box genes were named based on four ‘founding family members’, *<u>M</u>INICHROMOSOME MAINTENANCE 1 (MCM1)* from *Saccharomyces cerevisiae* (brewer’s yeast), *<u>A</u>GAMOUS* (*AG*) from *Arabidopsis thaliana* (*A. thaliana*; thale cress), *<u>D</u>EFICIENS* (*DEF*) from *Antirrhinum majus* (*A. majus*; snapdragon) and *<u>S</u>ERUM RESPONSE FACTOR* (*SRF*) from *Homo sapiens* (human) (Schwarz-Sommer et al., 1990). MADS-domain proteins bind as dimers to DNA-sequences including a 10-bp-long DNA element termed CArG-box (for C-A-rich-G), such as 5’-CC(A/T)_6_GG-3’ (Käppel et al., 2023).

Numerous whole genome analyses indicate that MADS-box genes exist only in eukaryotes. There is evidence, however, that the MADS-box originated from a DNA sequence encoding a region of subunit A of topoisomerase IIA in the stem group of extant eukaryotes (Gramzow et al., 2010). Even before the diversification of the crown group of extant eukaryotes had started, a gene duplication generated two lineages of MADS-box genes, termed Type I and Type II genes (Alvarez-Buylla et al., 2000; Gramzow and Theißen, 2010). The Type I and Type II genes of animals and humans are also known as *SERUM RESPONSE FACTOR-*like (*SRF*-like) and *MYOCYTE ENHANCER FACTOR 2*-like (*MEF2*-like) genes, respectively.

In all land plants (embryophytes) that have been investigated so far Type II genes have been annotated (Käppel et al., 2023). Typical Type II MADS-domain transcription factors of plants have a characteristic domain structure comprising four domains, the DNA-binding MADS-domain (M), the intervening domain (I), the keratin-like domain (K) and the variable C-terminal domain (C) (Kaufmann et al., 2005). These MADS-domain transcription factors were hence termed MIKC-type proteins, the corresponding genes MIKC-type genes. However, since not all Type II gene lineages have K boxes the terms ‘Type II genes of plants’ and ‘MIKC-type genes’ are not synonymous (Käppel et al., 2023), even though they are often used that way in the literature.

The K domain folds into coiled-coil domains involved in dimeric and tetrameric protein-protein interactions (Theißen et al., 2016; Käppel et al., 2023). These interactions underlie the versatile combinations of some MIKC-type proteins that are required for combinatorial functions and target site recognition. The capacity to combine and to constitute ‘Floral Quartet-like Complexes’ (FQCs) composed of four MIKC-type proteins binding as a tetramer to two sites (typically CArG-boxes) on target gene DNA, is of functional importance *in planta*, e.g. for the establishment of floral determinacy and organ identity in the angiosperm *Arabidopsis thaliana* (Hugouvieux et al., 2018; Hugouvieux et al, 2024) and, by inference, probably in essentially all flowering plants. FQC formation may have contributed to the fact that some MIKC-type proteins got involved in the control of many crucial developmental processes in flowering plants, including developmental phase changes and the control of organ identity (Gramzow and Theißen, 2010; Theißen et al., 2016).

The available evidence strongly suggests that MIKC-type genes originated in the stem group of extant streptophytes (charophyte algae + embryophytes), when a Type II MADS-box gene acquired a K box (Kaufmann et al., 2005; Nishiyama et al., 2018; Käppel et al., 2023). Whole genome analyses revealed that there are very few MIKC-type genes in extant charophyte species (Nishiyama et al, 2018; Gramzow et al., 2023; Feng et al., 2024), suggesting that there might have been just one MIKC-type gene in the most recent common ancestor (MRCA) of extant land plants. The ancestral MIKC-type gene present in charophytes was duplicated in the stem group of extant embryophytes (land plants), resulting in the lineages of MIKC^C^-type and MIKC*-type genes (Henschel et al., 2002; Gramzow and Theißen, 2010; Nishiyama et al., 2018; Rümpler et al., 2023).

The number of MIKC-type genes increased considerably during embryophyte evolution, often by the preferential retention of genes after whole genome duplications followed by sub- and neofunctionalization (Gramzow and Theißen, 2010; Theißen and Gramzow, 2016). For example, there are just two MIKC-type genes (one MIKC^C^-type and one MIKC*-type) in the liverwort *Marchantia polymorpha*, and 17 in the moss *Physcomitrium patens*, but roughly about 50 different genes in a typical flowering plant genome (Gramzow and Theißen, 2010; Thangavel and Nayar, 2018). This increase in gene number may have facilitated the evolution of body plan complexity in the sporophyte of land plants (Theißen et al., 2000).

In flowering plants MIKC^C^-type genes are typically involved in sporophyte ontogeny (Gramzow and Theißen, 2010). The most iconic function of MIKC^C^-type MADS-domain transcription factors is in the specification of the identity of floral organs, most importantly stamens and carpels (including ovules), but also different kinds of perianth organs such as sepals, petals or tepals, depending on the species (Theißen et al., 2016). The family of MIKC^C^-type genes includes 12 and 17 well-defined clades that had been established in the stem group of extant seed plants (spermatophytes) and flowering plants, respectively (Gramzow et al., 2014). Often members of the same clade share very similar and conserved functions in diverse developmental processes, such as the *DEF*- and *GLOBOSA*- (*GLO*-) like genes that specify petal and stamen identity, and the *AG*-like genes that specify stamen and carpel identity (Theißen et al., 1996, 2000, 2016; Gramzow and Theißen, 2010).

A comprehensive understanding of the evolution of MADS-box gene functions during evolution is hampered by the fact that in addition to flowering plants mutant phenotypes have been published for the moss *Physcomitrium patens* (Koshimizu et al., 2018), but not for any lycophyte, monilophyte (ferns and their allies such as horsetails (equisetophytes) and whisk ferns (psilophytes)), or gymnosperm. While orthology and expression patterns suggest that many MIKC^C^-type genes of gymnosperms have quite similar functions as their closest angiosperm relatives, such conclusions cannot be drawn for non-seed plants. This lack of insight hampers the understanding of major evolutionary innovations, such as ovules and seeds. Here, a better understanding of the reproductive biology, including MADS-box genes, of monilophytes would be helpful, as they represent the sister group of seed plants.

Even though monilophytes include the second most species-rich clade of vascular plants, i.e. ferns (with about 10,500 extant species) (Marchant et al., 2022), they have long been quite neglected in molecular biology, despite their unquestionable morphological diversity and ecological importance. Reasons that may have contributed to this neglect are the gigantic genomes, very high chromosome numbers, and lack of genetic transformation systems, to mention just a few.

Nevertheless, one fern species, *Ceratopteris richardii* (also known under the pet name “C-fern”), has been advertised as a model system already decades ago (Hickok et al., 1995; Eberle et al., 1995). It has been studied e.g. in terms of its life cycle and sex-determination mechanisms using classical mutagenesis (Hickok et al., 1995; Eberle et al., 1995; Banks, 1994, 1999), but progress was slow due to the lack of many tools that are available for flowering plant models such as *Arabidopsis thaliana*. With the sequencing of the whole genome (Marchant et al., 2022) and the establishment of an efficient transformation system (Plackett et al. 2014) the situation has considerably changed for the better, however. Among other approaches, the function of MADS-box genes in a fern could now be systematically investigated involving transgenic technology and reverse genetics.

The first MADS-box genes from ferns were identified (as cDNAs) from different *Ceratopteris* species already almost 30 years ago (Münster et al., 1997; Kofuji et al., 1997; Hasebe et al., 1998). These genes later all turned out to be MIKC^C^-type genes (a term that had not been established at the time yet). Compared to the typical organ identity genes of flowering plants most of these genes showed surprisingly broad expression patterns in both the gametophyte and the sporophyte (Münster et al., 1997; Kofuji et al., 1997; Hasebe et al., 1998). This observation led to the hypothesis that these fern MIKC-type genes have broader functions than those of the highly specialized floral homeotic genes of flowering plants; for example, like *MCM1* of baker’s yeast they might be involved in more general aspects of transcriptional control, cell differentiation or reproductive development (Theißen et al., 2000).

A good case in point is the MIKC^C^-type gene *Ceratopteris MADS3* (*CRM3*), also known as *CMADS6*, which was among the first MADS-box genes that was identified in *Ceratopteris* (Münster et al., 1997; Hasebe et al., 1998; Theißen et al., 2000). Stringent Southern blot analysis revealed that *CRM3* is a single copy gene in the genome of *C. richardii* (Münster et al., 1997), which was later corrobotated by whole genome sequence analysis (Marchant et al., 2022). However, *CRM3* has relatively closely related paralogs (subfamily members) in the *Ceratopteris* genome, such as *CRM9* (Theißen et al., 2000; Zhang et al., 2024). On the folllowing we will refer to the respective subfamily of genes as, *CRM3*-like genes‘.

A short peptide motif at the C-terminal end of CRM3-like proteins suggests a close relationship to class B (DEFICIENS-/GLOBOSA-like, also known as APETALA3-/PISTILLATA-like) and B_sister_ floral organ identity proteins of angiosperms (Kramer et al., 1998; Theißen et al., 2000). This relationship is weakly supported by phylogeny reconstructions (Münster et al., 1997; One Thousand Plant Transcriptomes Initiative, 2019), but certainly warrants further investigations.

Initial expression studies by Northern hybridization revealed that *CRM3* is expressed in both the gametophytic and the sporophytic phase of the life cycle of *C. richardii* (Münster et al., 1997), even though some authors reported that the expression is restricted to the gametophytic phase (Hasebe et al., 1998). More detailed analyses corroborated expression of *CRM3* in both phases and at all stages of the fern life cycle (Di Rosa, 1998). Virtual northern blot analyses revealed that *CRM3* mRNA accumulates relatively strongly in the gametophyte and young sprorophyte, but vanishes in fertile fronds (Fig. 1). This expression pattern is remarkably different from that of the closely related *CRM9*, which showed a relatively strong expression only in juvenile sporophytes, and from the more distantly related *CRM6* (Fig. 1; Di Rosa, 1998), which belongs to a different subfamily of fern MADS-box genes (Münster et al., 1997; One Thousand Plant Transcriptomes Initiative, 2019; Zhang et al., 2024).

**Figure 1.**
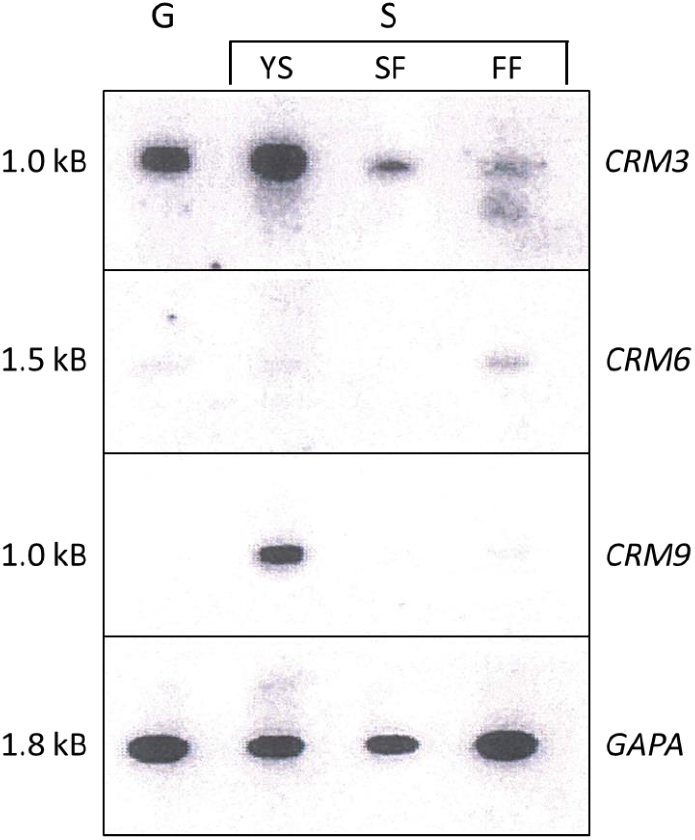
Virtual northern blot analysis of MADS-box gene expression in *C. richardii*. cDNAs were obtained from RNA harvested from the following material, as indicated above the lanes: G, Gametophyte (male and hermaphroditic); YS, young sporophyte (with 5 leaves); SF, sterile frond (without sporangia); FF, fertile frond (with sporangia); S, sporophyte. *CRM3, CRM6* and *CRM9* represent MADS-box genes. *GAPA* refers to a nuclear-encoded *Ceratopteris* glyceraldehyde-3-phosphate dehydrogenase used as loading control (Münster et al., 1997). (Figure taken from Di Rosa, 1998; modified.)

In situ hybridization studies suggested that in male gametophytes *CRM3* is expressed in spermatides, and in hermaphrodites in meristematic cells (Di Rosa, 1998; Theißen et al., 2000). In sporophytes *CRM3* as well as *CRM9* expression was found in the shoot axis as well as fronds of juvenile plants (Di Rosa, 1998; Theißen et al., 2000). In cross-sections of fertile fronds mRNAs of *CRM3, CRM6* and *CRM9* revealed different expression pattern, with especially *CRM6*, but also *CRM3* showing a relatively high mRNA accumulation in developing sporangia (Fig. 2; Di Rosa, 1998).

**Figure 2.**
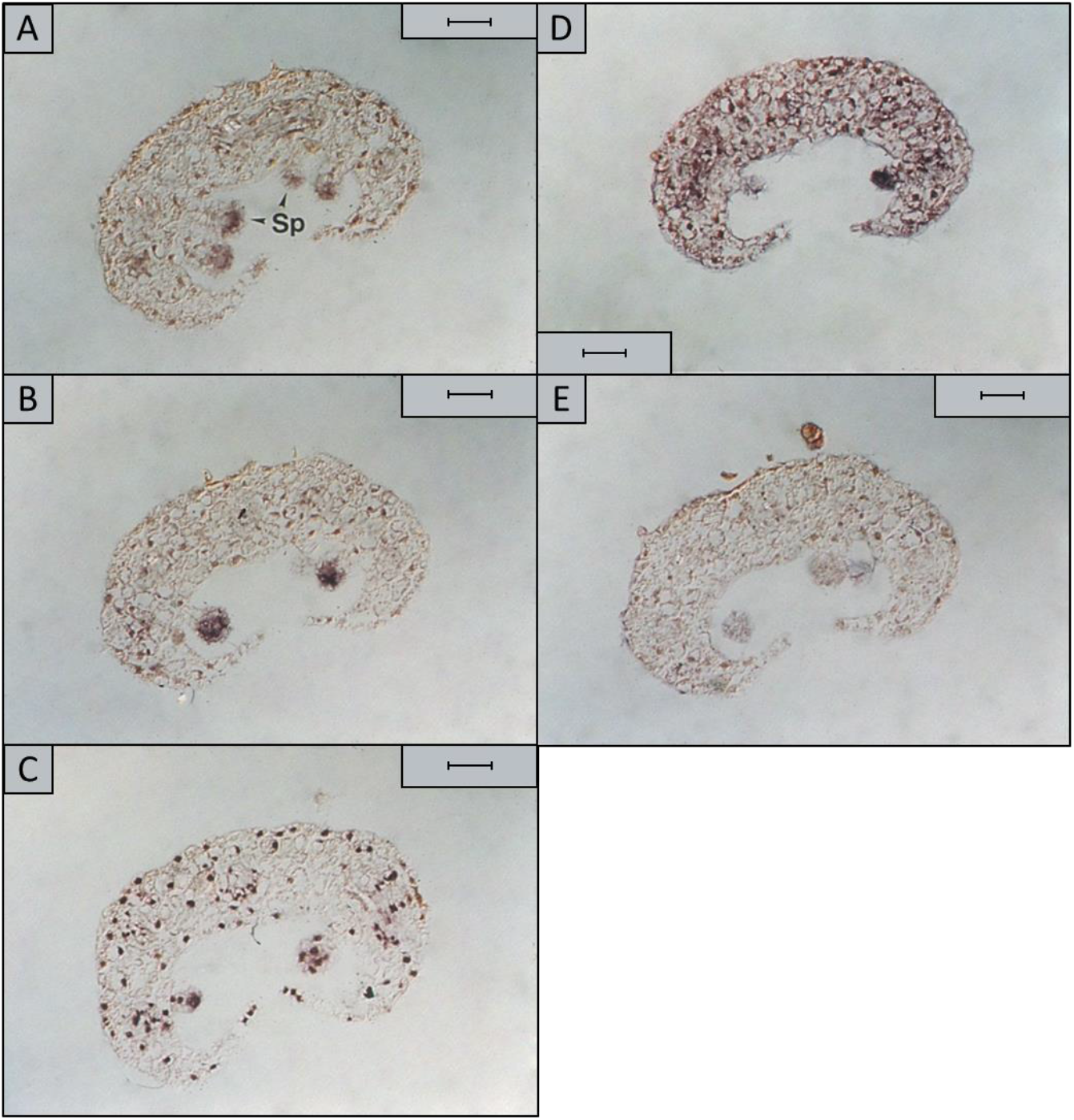
Expression of MADS-box genes in fertile fronds of *C. richardii* as revealed *by in situ* hybridization. The figure shows consecutive cross-sections of 7 μm thickness through a fertile frond at a plant age of 5 months. Expression is shown of (A*) CRM3*, (B) *CRM6*, (C) *CRM9*, (D) *GAPA*, as revealed by anstisense probes. In (E) hybridization was with a *CRM3* sense probe as control. Length of the bar in (A) – (E) represents 50 μm. sp, sporangia. (Figure taken from Di Rosa, 1998; modified.)

Despite these rather detailed insights into the expression pattern of *CRM3*, its function, like that of any other fern MADS-box gene, has remained elusive due to a lack of mutant phenotypes. As an initial attempt to learn more about *CRM3* function we generated transgenic plants that overexpress *CRM3* under control of the strong 35S promoter of the Cauliflower Mosaic Virus (CaMV).

## Materials and methods

### Plant materials and growth conditions

All work was done using *Ceratopteris richardii* strain Hn-n (Hickok et al., 1995). Plant growth conditions and media were as generally described (Plackett et al., 2014). The plants on media were grown in a phytochamber (Percival Scientific SE59-AR3, Perry, Iowa, USA) at 28 °C, 95 % humidity, 16 h of light / 8 h of dark with photosynthetic photon flux density (PPFD) of 180 µmol m^-2^ s^-1^. Since it turned out to be difficult to realize the high temperature and high humidity conditions optimal for *Ceratopteris* growth in our climate chamber we grew plants on soil in large transparent containers (plastic boxes; Böttcher, Sunware Q-Line Box 83300609) that were placed in a greenhouse (Fig. 3). The growth conditions inside the plastic boxes were 30 to 36 °C, > 90 % humidity, 16 h of light / 8 h of dark under two High-pressure sodium lamps. The spores were sterilised in 12 % [v/v] NaClO (Roth, Germany) for 8 min, washed five times and submerged in sterile water. After three days the spores were placed on C-fern media. In case of contamination 100 mg/L cefotaxime (ThermoFisher, Germany) was added to the media. All plants on soil were fertilized every two months with 200 mg/L (15-10-15) Planta Ferty 3 (PLANTA, Germany).

**Figure 3.**
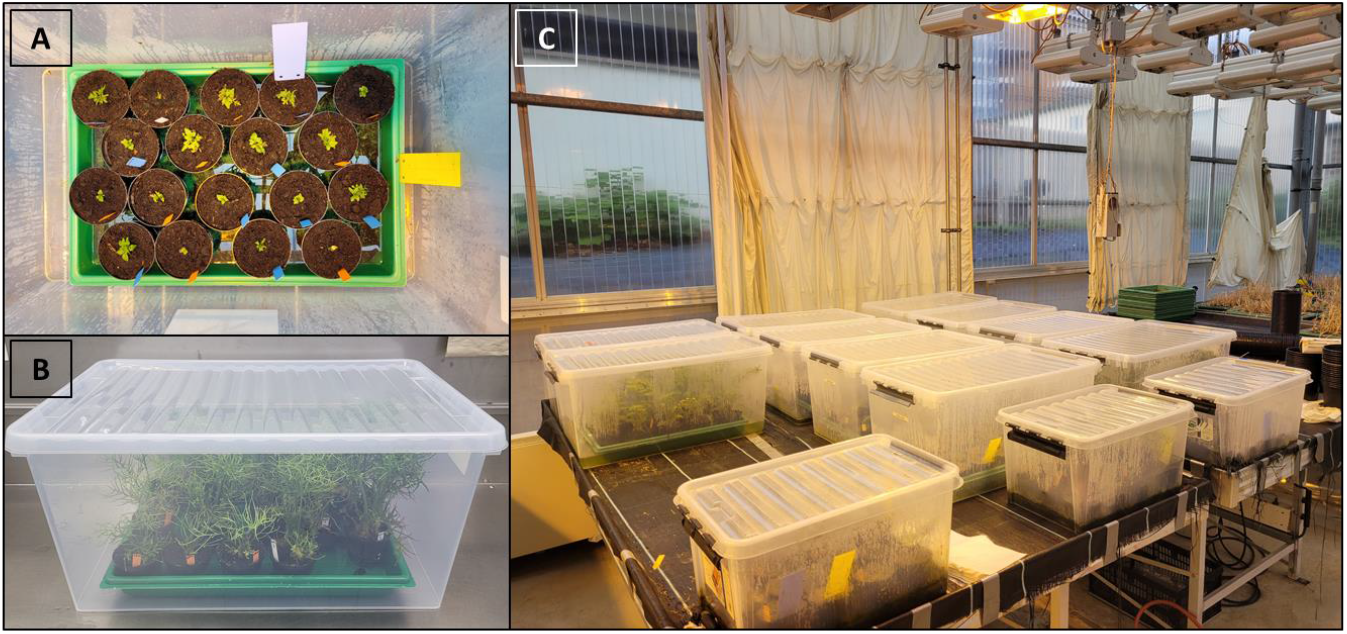
Conditions of fern growth. The transparent containers allow for a steady humidity of over 90%. When used in the greenhouse together with other cultures that need lower temperatures the containers also act as small greenhouses which show increased temperatures when directly under lamps. The left pictures (A and B) show transgenic 35S:*CRM3* and empty vector control 35S::*GUS* of *Ceratopteris richardii* 15 d and 72 d after moving to soil. The right picture (C) shows the setup of the containers inside the greenhouse.

### *Generation of constructs for overexpression of* CRM3

The construct pB-35S::*CRM3* was created by cloning the coding sequence (CDS) of *CRM3* (Phytozome: Ceric.15G037400.1) as a *Bam*HI-*Xba*I fragment into modified pBOMBER (Plackett et al., 2014). The recombination sites for *Bam*HI and *Xba*I were added to the CDS of *CRM3* by using *ECO31I* (FastDigest, ThermoFisher, Germany). Mutagenic primers were forward: 5’-GGTCTCAGATCATGGGCCGTGGAAAGATC-3’; reverse: 5’-GGTCTCGCTAGTTAGTTCAGTCTCAAGTCGAGGAAG-3’. A polymerase chain reaction (PCR) was done using these primers and Phusion polymerase (ThermoFisher, Germany). The PCR program was first 10 cycles with annealing at 63 °C for 15 s and elongation at 72 °C for 30 s, and then 20 cycles with the annealing changed to 72 °C (two step). The *GUS* gene of the original pBOMBER plasmid was cut out using *Bam*HI-*Xba*I (FastDigest, ThermoFisher, Germany) and the linear plasmid was isolated by gel purification (GeneJET Gel Extraction Kit, ThermoFisher, Germany). The mutagenic PCR product of *CRM3* was digested with *ECO31I* and ligated into the *Bam*HI-*Xba*I overhangs of the linear pBOMBER using T4-DNA ligase (ThermoFisher, Germany). The construct uses the CaMV 35S promoter for the expression of *CRM3* with OCS terminator. The selection is controlled by *hygromycin phosphotransferase* (*hpt*) for plant selection on hygromycin B.

### Generation of transgenic lines

Generation of callus, transformation and regeneration of callus into sporophytes was performed under aseptic conditions. Transformation was done by microparticle bombardment of callus essentially as described by Plackett et al. (2014). Exceptions are that only callus freshly grown from sporophytes was used for transformation and that the callus was selected on 25 mg/L hygromycin (ThermoFisher, Germany). Regeneration of sporophytes took 6 to 10 weeks. Once the sporophyte reached 1 to 2 cm in size and developed roots it was transferred on soil. First spores were collected 6-8 weeks after transferring the sporophyte to soil. Transgenic spores were selected on 5 mg/L hygromycin.

### Extraction of total genomic DNA

Total genomic DNA was extracted using a modified cetyltrimethylammonium bromide (CTAB) method with phenol-chloroform. 200 mg freshly frozen fronds (juvenile and fertile) were ground to dust with liquid nitrogen. 1 mL 2 % [w/v] CTAB pH 8.0 (Roth, Germany) combined with 10 µL 100 % ß-mercaptoethanol (Roth, Germany) were heated at 65 °C for 20 min and added to the frozen sample, homogenized by shaking and incubated at 65°C for 20 min. After centrifugation for 30 min at 16,000 g and 16°C the upper phase was taken und mixed with 500 µL phenol-chloroform [1:1] (Roth, Germany) and centrifuged for 30 min at 16,000 g and 16°C. The upper phase was taken and mixed with 0.1 volume 3 M [w/v] NaOAc pH 5.2 (Roth, Germany) and 0.7 volume [v/v] 100 % isopropanol (Roth, Germany). After incubation at room temperature for 60 min centrifugation for 1 h at 16.000 g and 4 °C the liquid was removed with a pipette and the pellet washed with 800 µL 70 % [v/v] ice-cold ethanol (Roth, Germany). The tube was incubated at 4 °C for 15 min, then centrifuged for 15 min. at 16,000 g and 4 °C. All liquid was removed carefully with a pipette and the pellet was dried on the bench at room temperature for 1 h. The DNA was dissolved by adding 50 µL Tris-EDTA (TE-buffer) pH 8.0 supplemented with 20 µg/µL RNase A (ThermoFisher, Germany) and incubated at 65 °C for 1 h, then stored at 4 °C.

### Extraction of total RNA

Total RNA was extracted from freshly frozen fronds (juvenile and fertile) using the RNeasy Kit, Mini, as described by the supplier (Qiagen, Germany). The first centrifugation step was repeated until all visible cell debris from the sample had been removed. The RNA pellet was dissolved in 40 µL nuclease free water.

### Inverse-PCR for number and location of transgenic insertion

Independent mutants and copy number of transgenic *CRM3* (35S::*CRM3*) were identified using inverse-PCR. 10 µg of total genomic DNA was digested in a volume of 50 µL with *Bam*HI (FastDigest, ThemoFisher, Germany) for 10 h at 37 °C. *Bam*HI was heat inactivated at 80 °C for 10 min. 50 µL of ligation mix using 15 U/µL T4-DNA ligase (ThermoFisher, Germany) supplemented with 1 mM ATP was directly added to the digestion mix and incubated for 12 h at 14 °C. The inverse PCR was done in two consecutive PCR reactions. The first PCR reaction involved 35 cycles in 20 µL volume with Phusion polymerase (ThermoFisher, Germany). The template was 2 µL unpurified product of cyclized genomic DNA fragments of the previous step. The primer pairs used for the first inverse PCR were P1 (forward: 5’-ATTAATTCAGTACATTAAAAACGTCCGC-3’; reverse: 5’-CACAACATACGAGCCGGAAG-3’) and located at the borders of the transgenic cassette. The second PCR involved 30 cycles in 20 µL volume with Phusion polymerase. The template was provided by using 2 µL directly from the previous PCR. The primer pair P2 (forward: 5’-GATCAAATATCATCTCCCTCGCAG, reverse GACGCTGTCGAAAATCGTGATC-3’) was located more outward of the transgenic cassette. The PCR program for both reactions was 98 °C for 2 min., then 35 cycles for P1 and 30 cycles for P2 with denaturation of 98 °C for 15 s, annealing of 64.5 °C for 30 s and elongation for 30 s followed by 72 °C for 10 min.

### Phenotypic characterization of overexpression mutants

Plants of the T_0_ generation of the wild-type, empty vector control 35S::GUS and 35S::CRM3 mutant were phenotyped once spores became visible on the plants which was at 60 to 80 d after transfering to soil. All fronds containing spores were cut and the total length of the fronds was measured. The mean values per plant were visualized in a violin plot using R (r-project.org) and statistical significance was calculated using Kruskal-Wallis multiple comparison and p-values adjusted with Bonferroni method in R using the packages FSA and dplyr.

### RT-qPCR to measure relative gene expression strength

Gene expression levels were identified using RT-qPCR. 500 ng of total RNA was used as template in cDNA synthesis, using Maxima Reverse Transcriptase (ThermoFisher, Germany). All primers were designed using the tool primer3 (https://primer3.ut.ee/). A list of primers is given in Table 1. The forward primer for the transgenic *CMR3* is located on the 5’-UTR following the CaMV 35S promoter. The forward primer for the endogenous *CRM3* is located on the 5’-UTR of the endogenous transcript. The reverse primer for both transgenic and endogenous *CRM3* is located on the CDS of exon 1. Specificity of primers was tested by sequencing the PCR products. RT-qPCR was done with three technical replicates. Each reaction was performed in a qPCR cycler Mx3005P (STRATAGENE, USA) with Maxima SYBR Green/ROX qPCR Master Mix (ThermoFisher, Germany). Primer efficiency was tested with a serial dilution of cDNA and calculated by (〖10〗 ^(−1/*slope*) − 1) * 100 in Excel 2019 (Microsoft). Expression of transgenic and endogenous *CRM3* was normalized against the expression of two housekeeping genes *CrACTIN-7* (Phytozome: Ceric.08G028400) and *CrTATA-BINDING PROTEIN* (*CrTBP*) (Phytozome: Ceric.29G071900). Relative expression was calculated by 2^-ΔCt^ method. The RT-qPCR mix per reaction was 20 µL containing 10 µL master mix, 0.6 µM final concentration per primer and 1 µL of 1:5 diluted cDNA. The program was 95 °C for 10 min., then 40 cycles of 95 °C for 15 s, 60 °C for 30 s and 72 °C for 30 s.

**Table 1:**
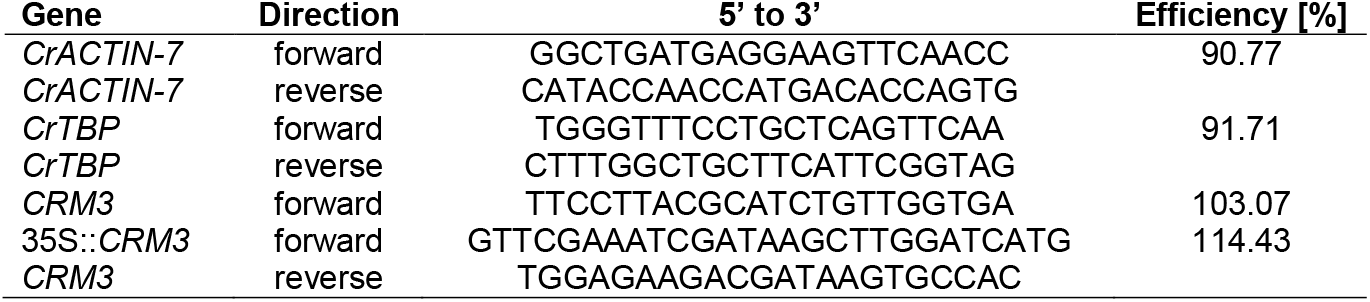
Primers used for RT-qPCR on cDNA of transgenic *CRM3* overexpression mutants. The reverse primer for *CRM3* was used for both endogenous and transgenic *CRM3*.

## Results

In order to overexpress cDNA of the MIKC^C^-type MADS-box gene *CRM3* in *C. richardii*, plasmids with a gene cassette containing *CRM3* cDNA under control of the CaMV 35S promoter (pB-35S::*CRM3*) were generated and transformed into *C. richardii*. After transforming the callus by microparticle bombartment, it was selected on antibiotic media for six weeks which involves transferring the callus to fresh media every two weeks, separating green callus from dead callus. After four to six weeks the callus started to differentiates into sporophytes. During that process it often happened that the callus broke into smaller pieces or that multiple sporophytes grew out of a single piece of callus. It was, therefore, tested whether regenerated sporophytes are clones originating from the same cell of callus or independent mutants. For this purpose, inverse PCR based on genomic DNA (gDNA) was performed.

First, putative mutants were genotyped by PCR on gDNA extracted of juvenile fronds 14 d after transferring the plants to soil (Fig. 4A). PCR to test for the presence of the full *CRM3* transgene confirmed about 48 % (n = 98) of the transformed plants to be positive. The four mutants 35S::*CRM3*_1, _2, _3 and _9 are used here as examples. They were tested positive among the first ten plant being transferred to soil. We confirmed that *CRM3* was expressed under the 35S promoter by a PCR on cDNA with primers binding specifically to the transgenic UTR regions (Fig. 4B). To test if the mutants are independent or clones the previously isolated gDNA was digested with *BamH*I (Fig. 4C) and the fragments were cyclized due to the self-compatible overhangs (Fig 4D). A PCR in which the primers are facing outwards while located on the transgene has amplified the flanking regions upstream and downstream to the transgene until the closest *BamH*I site on the gDNA. All tested mutants show a unique band pattern between one and nine bands which confirms that they are indeed independent mutants containing one to several copies of the transgene (Fig. 4E). The wild-type shows no PCR product or artefacts, which confirms the methodology (Fig. 4E).

**Figure 4.**
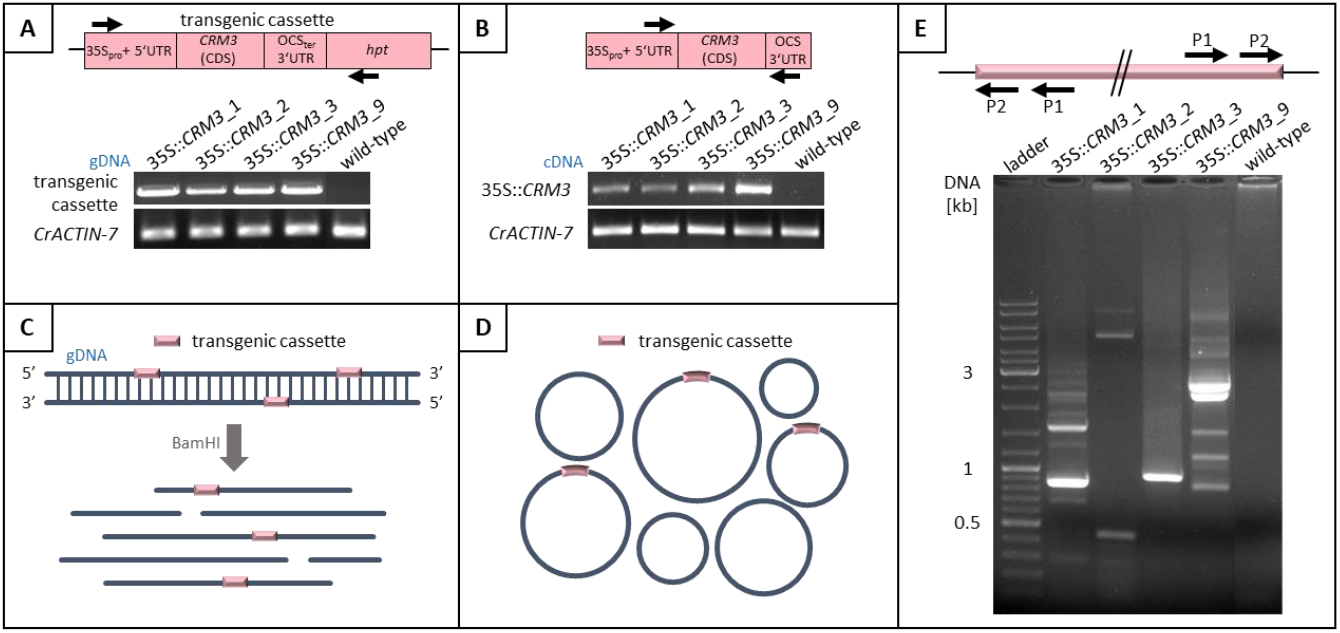
Presence, copy number and expression of a *CRM3* cassette in transgenic *C. richardii*. (A) Genotyping: PCR on genomic DNA (gDNA) revealing the presence of transgenic cassette including CaMV 35S promoter with synthetic 5’-UTR, coding region of *CRM3*, OCS terminator and *hpt* gene for hygromycin resistance. (B) Test for expression of transgenic *CMR3*. The primers are located on the transgenic UTR regions to exclude mRNA of endogenous *CRM3*. (C) – (E) Procedure for reverse PCR. (C) Scheme for restriction digestion of gDNA. (D) Scheme depicting how fragments obtained in (C) were cyclized with T4-ligase. (E) The flanking regions of the transgene were amplified using outward facing primers and PCR products were separated on an agarose gel.

According to our experience, regenerating the sporophyte of *C. richardii* from callus causes always a disturbed phenotype. This effect becomes much less prominent with age of the plant. It has previously been shown that *CRM3* is expressed throughout the entire life-cycle of the sporophyte (Fig. 1; Münster et al., 1997; Di Rosa, 1998) and that it hence could be involved in spore development. In an initial study on the phenotypic consequences of *CRM3* overexpression we, therefore, focused on the sporophyll.

The reproductive phase of the life cycle of the sporophyte of *Ceratopteris richardii*, in which every new grown frond contains spores, is reached at 60 to 80 days after transferring to soil (120 to 150 days after transformation). After confirming the presence of spores inside a frond it was cut and the length of the frond was measured. It turned out that the sporophylls of T_0_ 35S::*CRM3* transgenics (n = 48) (e.g. Fig. 5B, E) had a mean length of 12.4 cm, whereas those of the empty vector control 35S::*GUS* (n = 8, e.g. Fig. 5D) had a mean length of 23.1 cm and the wild-type (n = 52, e.g. Fig. 5A, C) a mean of 19.5 cm. Thus 35S::*CRM3* transgenics had an on average significantly shorter sporophylls and also a much wider distribution of sporophyll length (Fig. 5F). The empty vector control did not show a significant difference to the wild-type. Both 35S::*GUS* and wild-type had been regenerated from callus in parallel to 35S::*CRM3* plants.

**Figure 5:**
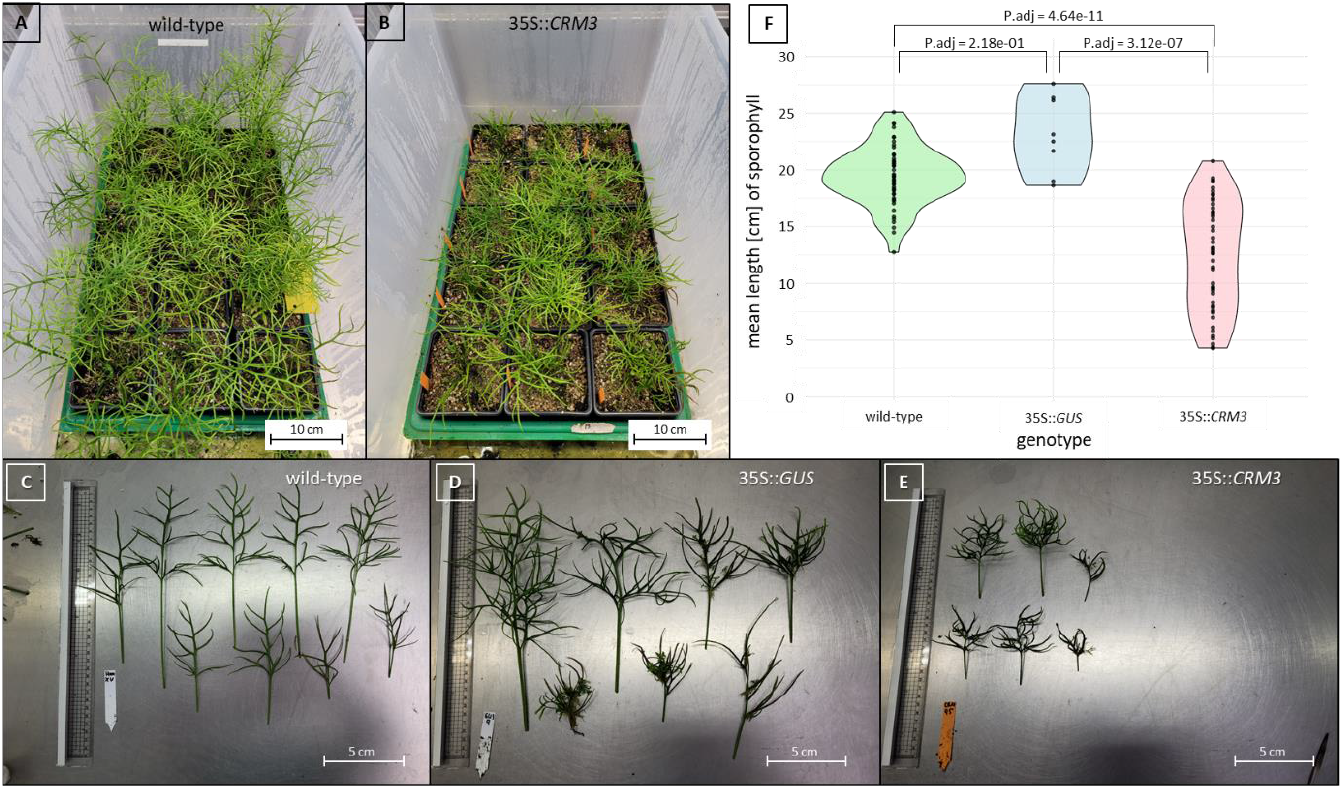
Phenotyping of the sporophylls of adult *C. richardii* plants. (A) Wild-type plants (n = 15, 80 days on soil) with direct comparison to (B) *CRM3* overexpression mutants 35S::*CRM3* (n = 15, 80 days on soil). The fronds containing spores of (C) wild-type (mean = 19.5 cm), (D) empty vector control 35S::*GUS* (mean = 19.0 cm) and (E) 35S::*CRM3* (mean = 12.0 cm) were cut and measured. (F) Distribution of mean length of sporophylls of the genotypes wild-type, 35S::*GUS* and 35S::*CRM3*; statistical significance was calculated using Kruskal-Wallis multiple comparison and p-values adjusted with Bonferroni method in R (r-project.org) using the packages FSA and dplyr.

RT-qPCR (Fig. 6J) on total RNA as template was performed to understand the expression strength of 35S::*CRM3* in relation to endogenous *CRM3* and how it changes between the juvenile (Fig. 6G) and reproductive (Fig. 6H) stage of *Ceratopteris richardii*. The primer pairs for endogenous *CRM3* and 35S::*CRM3* bind to the respective 5’-UTR and the first exon of the CDS of *CRM3* (Fig. 6I). Total RNA from the mutants (transgenics) 35S::*CRM3*_1, _2, _3 and _9 (Fig. 6A-D) was isolated after 14 days on soil representing the early juvenile stage and 120 days on soil representing the late reproductive stage. The C_t_ values of endogenous *CRM3* and 35S::*CRM3* were normalized against the housekeeping genes *CrACTIN-7* and *CrTBP*. The expression strength was calculated with the 2^-ΔCt^ method. All expression values were divided by the value of endogenous *CRM3* in the juvenile stage to show the difference in expression between the juvenile and reproductive stage. All four mutants tested showed that the expression strength of endogenous *CRM3* strongly decreases in the reproductive stage. It also shows that the expression strength of 35S::*CRM3* varies highly between the tested mutant but also remains steady between juvenile and reproductive stage. The mutant 35S::*CRM3*_9 showed the highest overexpression in comparison to endogenous *CRM3* with a fold of 5.7 in juvenile tissue and 35.8 in reproductive tissue. The inverse PCR of mutant 9 showed many bands which suggested a high copy number of 35S::*CRM3* (Fig. 4E). In contrast to this the mutant 35S::*CRM3*_1 showed a high amount of bands as well but the RT-qPCR only showed an overexpression of 35S::*CRM3* of 0.3 fold in juvenile tissue and 10.1 fold in reproductive tissue.

**Figure 6:**
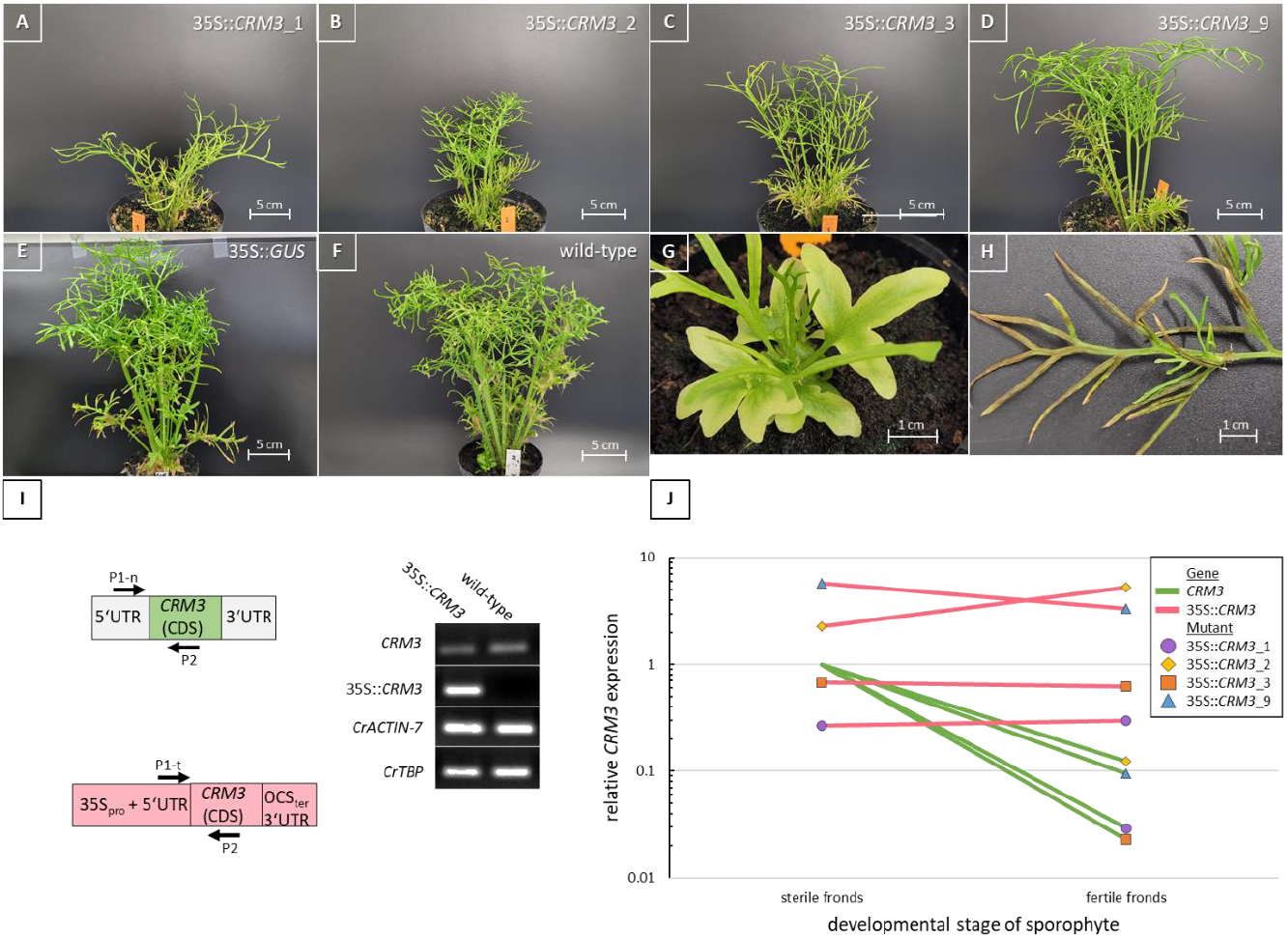
Relative overexpression of 35S::*CRM3* in comparison to endogenous *CRM3*. (A-D) Phenotypes of the T_0_ 35S::*CRM3* mutants 1, 2, 3 and 9 in comparison to (E) empty vector control 35S::*GUS* and the (F) non-transgenic wild-type. Total RNA was isolated from 5 different (G) juvenile and (H) reproductive fronds. (I) Primer pairs for RT-qPCR was designed so that the forward primer was specific to endogenous (P1-n) or transgenic (P1-t) *CRM3* while the reverse primer was located in the coding region of the first exon of *CRM3*. (J) RT-qPCR results of the four mutants with the expression of 35S::*CRM3* in relation to endogenous *CRM3* shown on a log(10) scale. The mean Ct values were first normalized against the housekeeping genes *CrActin-7* and *CrTBP*. The expression was calculated with the 2^-ΔCt^ method. All expression values were divided by endogenous *CRM3* in the juvenile stage to show the difference between the juvenile and mature stage.

The average length of the mature sporophylls of the mutants are 13.6 cm for 35S::*CRM3*_1, 10.0 cm for 35S::*CRM3*_2, 11.4 cm for 35S::*CRM3*_3, and 19 cm for 35S::*CRM3*_9 which is closed to the wild-type with an average of 19.5 cm (n = 52) while the other three are about 9 cm smaller (Fig. 5).

## Discussion

We report here, to the best of our knowledge, the first overexpression of any MADS-box gene in a fern, exemplified with *CRM3* in *Ceratopteris richardii*. Clear phenotypic differences are visible of *CRM3* overexpressing plants compared to wild-type and empty-vector control plants, suggesting that the observed effects on sporophyll development are not simply due to the origin of the plants from callus or the transformation procedure.

Generations beyond T_0_ will be analyzed in future investigations in order to determine whether the transgenic phenotype is stably inherited or subject to phenomena such as gene silencing. Due to two endogenous wild-type alleles of *CRM3* in addition to the transgenes in the *Ceratopteris richardii* genome this is a possible fate of both the artificially 35S promoter-driven transgenic and the endogenous copies that cannot be completely excluded.

In addition to the sporophytic generation, also gametophytes (both hermaphroditic and male), i.e. the whole life cycle of *Ceratopteris richardii* need to be analyzed in the future to get a comprehensive overview about overexpression effects of *CRM3*. The life-cycle has been documented for the wild-type in great detail already (Conway and Di Stilio, 2019), which will be very helpful for recognizing differences brought about by transgene overexpression. Comparative studies involving related MIKC^C^-type MADS-box genes such as *CRM6* and *CRM9* might also be revealing by indicating to what extent the phenotypic effects seen are gene specific.

As a means to get clues about unknown gene functions the ectopic and overexpression of genes, often even used in a heterologous background, has been applied many times in plant research. Often the CaMV 35S promoter was used in such experiments to drive strong, almost ubiquitous transgene expression. Sometimes the overexpression phenotypes are highly informative, e.g. in case of organ identity (homeotic) genes. For example, ectopic expression of endogenous *AP3* and *PI* under control of the 35S promoter in floral primoria of *Arabidopsis thaliana* transforms sepal primordia into petals and carpel primordia into stamens, indicating that *AP3* and *PI* function as class B floral organ identity genes specifying petal and stamen identity (Riechmann and Meyerowitz, 1997). This conclusion is corroborated by the quite specific expression pattern of *AP3* and *PI* in petal and stamen primordia in the wild-type, and by the loss-of-function phenotype of these genes (development of sepals and carpels rather than petals and stamens, respectively).

In many other cases, however, results of overexpression experiments are not so easy to interpret, the approach has hence clear limitations. Expression of a gene at a wrong place (tissue, organ), a wrong time or at an unphysiologically high level, may just bring about quite unspecific effects that have little to do with the gene function proper. For example, the ectopic expression of many genes in plants, including MADS-box genes, causes early flowering, as has been shown in literally hundreds (if not thousands) of studies (e.g., Yu et al., 2025). Whether this phenotype reveals a flowering time gene proper or is just due to an indirect (e.g. stress) response needs to be determined in every single case (but as a matter fact, often isn’t).

The possibility of an unspecific effect currently cannot be completely excluded for *CRM3*. In contrast to typical floral homeotic genes such as *AP3* and *PI, CRM3* shows in the wild-type a temporally and spatially much broader expression pattern that is more difficult to correlate with any transgenic phenopyte and wild-type gene function. To what extent *CRM3* functions in cell proliferation and elongation, for example, as suggested by the phenopyte of transgenic sporophytes, remains to be seen in future investigations.

More insights into *CRM3* function are expected from the study of knock-out plants displaying loss-of-function phenotypes. The generation of such mutants employing the CRISPR-Cas9 system is well underway.

## Acknowledgements

We are grateful to Andrew Placket for his valuable help with the establishment of the *Ceratopteris* transformation system. Many thanks to Annette Becker for numerous valuable discussions on *Ceratopteris* research over three decades. In addition we also thank all other members of the ICIPS research unit for fruitful discussions.

## References

Alvarez-Buylla ER, Pelaz, S, Liljegren SJ, Gold SE, Burgeff C, Ditta GS, de Pouplana LR, Martínez-Castilla L, Yanofsk MF (2000) An ancestral MADS-box gene duplication occurred before the divergence of plants and animals. Proc. Natl. Acad. Sci. USA 97:5328–5333. DOI: 10.1073/pnas.97.10.5328

Banks JA (1994) Sex-determining genes in the homosporous fern Ceratopteris. Development 120(7):1949–1958. DOI: 10.1242/dev.120.7.1949

Banks JA (1999) Gametophyte development in ferns. Annu. Rev. Plant Physiol. Plant Mol. Biol. 50:163–186. DOI: 10.1146/annurev.arplant.50.1.163

Conway SJ, Di Stilio VS (2019) An ontogenetic framework for functional studies in the model fern Ceratopteris richardii. Dev. Biol. 457:20–29. DOI: 10.1016/j.ydbio.2019.08.017

Di Rosa A (1998) Molekularbiologische Untersuchungen zum Ursprung homöotischer Gene in Pflanzen am Beispiel der MADS-Box-Genfamilie aus dem Farn Ceratopteris richardii. Dissertation, Mathematisch-Naturwissenschaftliche Fakultät der Universität zu Köln.

Eberle J, Hasebe M, Nemancheck J, Wen C-K, Banks JA (1995) Ceratopteris: a model system for studying sex-determining mechanisms in plants. Int. J. Plant Sci. 156:359–366. DOI: 10.1086/297257

Feng X, Zheng J, Irisarri I, et al. (2024) Genomes of multicellular algal sisters to land plants illuminate signaling network evolution. Nature Genetics 56:1018–1031. DOI: 10.1038/s41588-024-01737-3

Gramzow L, Theißen G (2010) A hitchhiker’s guide to the MADS world of plants. Genome Biol. 11:214. DOI: 10.1186/gb-2010-11-6-214

Gramzow L, Ritz MS, Theißen G (2010) On the origin of MADS-domain transcription factors. Trends Genet. 26:149–153. DOI: 10.1016/j.tig.2010.01.004

Gramzow L, Tessari C, Rümpler F, Theißen G (2023) Deep evolution of MADS-box genes in Archaeplastida. bioRxiv 10.1101/2023.02.13.528266

Gramzow L, Weilandt L, Theißen G (2014) MADS goes genomic in conifers: Towards determining the ancestral set of MADS-box genes in seed plants. Ann. Bot. 114:1407–1429. DOI: 10.1093/aob/mcu066

Hasebe M, Wen C-K, Kato M, Banks JA (1998) Characterization of MADS homeotic genes in the fern Ceratopteris richardii. Proc. Natl. Acad. Sci. USA 95:6222–6227. DOI: 10.1073/pnas.95.11.6222

Henschel K, Kofuji R, Hasebe M, Saedler H, Münster T, Theißen G (2002) Two ancient classes of MIKC-type MADS-box genes are present in the moss Physcomitrella patens. Mol. Biol. Evol. 19:801–814. DOI: 10.1093/oxfordjournals.molbev.a004137

Hickok LG, Warne TH, Fribourg RS (1995) The biology of the fern Ceratopteris and its use as a model system. Int. J. Plant Sci. 156:332–345. DOI: 10.1086/297255

Hugouvieux V, Silva CS, Jourdain A, Stigliani A, Charras Q, Conn V, Conn SJ, Carles CC, Parcy F, Zubieta C (2018) Tetramerization of MADS family transcription factors SEPALLATA3 and AGAMOUS is required for floral meristem determinacy in Arabidopsis. Nucleic Acids Res. 46:4966–4977. DOI: 10.1093/nar/gky205

Hugouvieux V, Blanc-Mathieu R, Janeau A, Paul M, Lucas J, Xu X, Ye H, Lai X, Le Hir S, Guillotin A, Galien A, Yan W, Nanao M, Kaufmann K, Parcy F, Zubieta Ch (2024) SEPALLATA-driven MADS transcription factor tetramerization is required for inner whorl floral organ development. The Plant Cell 36:3435–3450. DOI: 10.1093/plcell/koae151

Käppel S, Rümpler F, Theißen G (2023) Cracking the floral quartet code: how do multimers of MIKC^C^-type MADS-domain transcription factors recognize their target genes? International Journal of Molecular Sciences 24:8253. DOI: 10.3390/ijms24098253

Kaufmann K, Melzer R, Theißen G (2005) MIKC-type MADS-domain proteins: Structural modularity, protein interactions and network evolution in land plants. Gene 347:183–198. DOI: 10.1038/nprot.2009.244

Kofuji R, Yamaguchi K (1997) Isolation and phylogenetic analysis of MADS genes from the fern Ceratopteris richardii. J Phytogeogr Taxon 45:83–91. 10.24517/00055551

Koshimizu S, Kofuji R, Sasaki-Sekimoto Y., et al. (2018) Physcomitrella MADS-box genes regulate water supply and sperm movement for fertilization. Nature Plants 4:36–45. 10.1038/s41477-017-0082-9

Kramer EM, Dorit RL, Irish VF (1998) Molecular evolution of genes controlling petal and stamen development: duplication and divergence within the APETALA3 and PISTILLATA MADS-box gene lineages. Genetics 149:765–783. DOI: 10.1093/genetics/149.2.765

Marchant DB, Chen G, Cai S, Chen F, et al. (2022) Dynamic genome evolution in a model fern. Nature Plants 8:1038–1051. 10.1038/s41477-022-01226-7

Münster T, Pahnke J, Di Rosa A., Kim JT, Martin W, Saedler H, Theißen G (1997) Floral homeotic genes were recruited from homologous MADS-box genes preexisting in the common ancestor of ferns and seed plants. Proc. Natl. Acad. Sci. USA 94:2415–2420. DOI: 10.1073/pnas.94.6.2415

Münster T, Faigl W, Saedler H, Theißen G (2002) Evolutionary aspects of MADS-box genes in the eusporangiate fern Ophioglossum. Plant Biol. 4:474–483. DOI: 10.1055/s-2002-34130

Nishiyama T, Sakayama H, de Vries J, Buschmann H, Saint-Marcoux D, Ullrich KK, Haas FB, Vanderstraeten L, Becker D, Lang D, et al. (2018) The Chara genome: secondary complexity and implications for plant terrestrialization. Cell 174:448–464.e24. DOI: 10.1016/j.cell.2018.06.033

One Thousand Plant Transcriptomes Initiative (2019) One thousand plant transcriptomes and phylogenomics of green plants. Nature 574:679–685. DOI: 10.1038/s41586-019-1693-2

Plackett ARG, Huang L, Sanders HL, Langdale JA (2014) High-efficiency stable transformation of the model fern species Ceratopteris richardii via microparticle bombardment. Plant Physiology 165:3–14. DOI: 10.1104/pp.113.231357

Riechmann JL, Meyerowitz EM (1997) MADS domain proteins in plant development. Biol. Chem. 378:1079-1101. PMID: 9372178

Rümpler F, Tessari C, Gramzow L, Gafert C, Blohs M, Theißen G (2023) The origin of floral quartet formation—Ancient exon duplications shaped the evolution of MIKC-type MADS-domain transcription factor interactions. Mol. Biol. Evol. 40:msad088. DOI: 10.1093/molbev/msad088

Schwarz-Sommer Z, Huijser P, Nacken W, Saedler H, Sommer H (1990) Genetic control of flower development by homeotic genes in Antirrhinum majus. Science 250:931–936. DOI: 10.1126/science.250.4983.931

Thangavel G, Nayar SA (2018) Survey of MIKC type MADS-box genes in non-seed plants: Algae, Bryophytes, Lycophytes and Ferns. Front. Plant Sci. 9:510. DOI: 10.3389/fpls.2018.00510

Theißen G, Becker A, Di Rosa A, Kanno A, Kim JT, Münster T, Winter K-U, Saedler H (2000) A short history of MADS-box genes in plants. Plant Molecular Biology 42:115–149. DOI: 10.1007/BF02337521

Theißen G, Gramzow L (2016) Structure and evolution of plant MADS-domain transcription factors; In Plant Transcription Factors: Evolutionary, Structural and Functional Aspects; Gonzalez, D.H., Ed.; Elsvier: Philadelphia, PA, USA, pp.127–138. DOI: 10.1016/B978-0-12-800854-6.00008-7

Theißen G, Kim J, Saedler H (1996) Classification and phylogeny of the MADS-box multigene family suggest defined roles of MADS-box gene subfamilies in the morphological evolution of eukaryotes. J. Mol. Evol. 43:484–516. DOI: 10.1007/BF02337521

Theißen G, Melzer R, Rümpler F (2016) MADS-domain transcription factors and the floral quartet model of flower development: Linking plant development and evolution. Development 143:3259–3271. DOI: 10.1242/dev.134080

Yu H, Xia L, Zhu J, Xie X, Wie Y, Li X, He X, Luo C (2025) Genome-wide analysis of the MADS-box gene family in mango and ectopic expression of MiMADS77 in Arabidopsis results in early flowering. Gene 935:149054. DOI: 10.1016/j.gene.2024.149054

Zhang R, Zhang J, Xu Y-X, Sun J-M, Dai S-J, Shen H, Yan Y-H (2024) Dynamic evolution of MADS-box genes in extant ferns via large-scale phylogenomic analysis. Frontiers in Plant Science 15:1410554. DOI: 10.3389/fpls.2024.1410554

